# Single-cell and spatiotemporal transcriptomic profiling of brain immune infiltration following Venezuelan equine encephalitis virus infection

**DOI:** 10.1101/2024.09.12.612602

**Authors:** Margarita V. Rangel, Aimy Sebastian, Nicole F. Leon, Ashlee M. Phillips, Bria M. Gorman, Nicholas R. Hum, Dina R. Weilhammer

## Abstract

Neurotropic alphaviruses such as Venezuelan equine encephalitis virus (VEEV) are critical human pathogens that continually expand to naïve populations and for which there are no licensed vaccines or therapeutics. VEEV is highly infectious via the aerosol route and is a recognized weaponizable biothreat that causes neurological disease in humans. The neuropathology of VEEV has been attributed to an inflammatory immune response in the brain yet the underlying mechanisms and specific immune cell populations involved are not fully elucidated. This study uses single-cell RNA sequencing to produce a comprehensive transcriptional profile of immune cells isolated from the brain over a time course of infection in a mouse model of VEEV. Analyses reveal differentially activated subpopulations of microglia, including a distinct type I interferon-expressing subpopulation. This is followed by the sequential infiltration of myeloid cells and cytotoxic lymphocytes, also comprising subpopulations with unique transcriptional signatures. We identify a subpopulation of myeloid cells that form a distinct localization pattern in the hippocampal region whereas lymphocytes are widely distributed, indicating differential modes of recruitment, including that to specific regions of the brain. Altogether, this study provides a high-resolution analysis of the immune response to VEEV in the brain and highlights potential avenues of investigation for therapeutics that target neuroinflammation in the brain.

**Author Summary:** Venezuelan equine encephalitis virus (VEEV) causes brain inflammation in both animals and humans when transmitted by mosquito bite or infectious aerosols. The mechanisms underlying disease caused by VEEV, including the role of the immune response in brain pathology, are not well understood. Here we performed a comprehensive assessment of the immune response to VEEV in the brain over time using two advanced sequencing techniques. Following infection, immune cells infiltrate the brain in a sequential fashion and display different activation profiles. Different types of immune cells also display strikingly different spatial patterns throughout the brain. This study provides the most comprehensive description of the immune response to VEEV in the brain performed to date and advances our understanding of immune-driven neuropathology and identification of therapeutic targets.

## Introduction

Venezuelan equine encephalitis virus (VEEV) is a mosquito-borne virus within the *Alphavirus* genus and *Togaviridae* family that has caused devastating outbreaks of disease among humans and equine in the Americas over recent decades [1–3]. VEEV is maintained in a sylvatic cycle between mosquitos and wild rodents but can spill over and amplify to high titers in equine, leading to epizootic and epidemic outbreaks. VEEV is a New World alphavirus, which are mostly encephalitic, as opposed to the Old World alphaviruses, which cause mostly arthritogenic disease. The virus causes an acute febrile illness that can lead to severe and lethal encephalitic cases with neurological symptoms spanning dizziness, headache, confusion, seizures, and stroke. While the mortality rate overall is <1% in adults and <5% in children, central nervous system (CNS) infections and neurological manifestations can occur in up to 14% of cases and among encephalitic cases, risk of mortality increases up to 10% in adults and 35% in children [4, 5]. In addition to the natural transmission route via mosquito bite, humans are also susceptible to infection via aerosol exposure, as demonstrated by the report of multiple lab-acquired infections [6, 7]. This, paired with the fact that VEEV can be grown to high titers, contributes to its designation as a category B priority pathogen by the National Institutes of Health and as a select agent by the Centers for Disease Control and Prevention. There are no specific treatments available for VEEV infection nor virus-induced encephalitis. A challenge in treating such infections is an incomplete understanding of the key drivers of pathogenesis, specifically the cellular and molecular changes underlying the progression of damaging inflammation that occurs in the brain. Thus, a temporal and spatial characterization of VEEV pathogenesis in the brain would elucidate the mechanisms of disease and the role of the host immune response, as well as aid the identification of potential therapeutic targets.

Previous studies have established that inflammation is a key component of VEEV neuropathogenesis. Severe vascular cuffing, cellular infiltration, edema, and neuronal karyorrhexis can be observed histologically in VEEV-infected brains [8–10]. An elevation of pro-inflammatory cytokines (e.g. IL-1β, IL-6, IL-12, TNF-α, and IFN-γ) and components of antigen presentation, apoptosis, and antiviral response (e.g. Cxcl9, Cxcl10, Cxcl11, Ccl2, Ccl5, Ifr7, Ifi27, Oas1b, Fcerg1, Mif, Clusterin, and MHC class II), have been detected in VEEV-infected mouse brains via histological, cytokine array, and RNA microarray techniques applied to brain homogenate or total RNA [11–14]. The host immune response plays paradoxical roles during VEEV infection in that a sufficiently robust response is required for systemic clearance, meanwhile cell depletion experiments and experiments in immune-compromised mice demonstrate a dampened response can improve outcome [15, 16]. These findings suggest that immune invasion in the brain is a large contributor to disease outcome. Thus, defining functions of discrete populations of immune cells that infiltrate the brain during infection would provide an opportunity to identify targetable pathways for immunomodulatory therapies.

There is currently an incomplete understanding of the specific activation states of resident and infiltrating immune cells and the roles they play during the inflammatory response to VEEV infections. Further, the kinetics of the response and sequential recruitment of these cells has not been thoroughly described. Here, we aimed to bridge this gap in knowledge by generating a high-resolution profiling of the transcriptional activity of individual immune cells in the brain over a time course of VEEV infection using an established murine model, paired with spatial transcriptomic analysis during severe neuropathology. We applied single-cell RNA sequencing (scRNAseq) to immune cells isolated from the brains of C3H/HeN mice infected with VEEV TC-83 at 2, 4, and 6 dpi. Fluctuations in the immune populations present in the brain were observed during these key timepoints of disease development. We identified sequentially emerging subpopulations of microglia and infiltrating myeloid and lymphoid cells with unique transcriptional profiles. Spatial transcriptomic analysis on brains at 6 dpi revealed differential distribution of immune cell subtypes, with a subtype of myeloid cells localizing at the periphery of the cortex and hippocampal region of infected brains whereas other myeloid and lymphoid groups were more widespread. These results comprise a comprehensive temporal profiling of transcriptomic activity in the brain in response to VEEV infection that provide insight into key immune players during the onset of neuroinflammation, information crucial to a fundamental understanding of VEEV pathogenesis and the potential design of host-based immunomodulatory therapeutics. Additionally, spatial characterization of key populations during a highly diseased state highlights specific regions of the brain exhibiting differential immune activity that may be important to consider for effective delivery of brain-targeting treatments.

## Results

### Single-cell RNA sequencing reveals robust and sequential immune cell infiltration in mouse brains during VEEV infection

To understand the progression of the neuroinflammatory immune response to VEEV, we profiled the fluctuations of immune cell populations in the brain and their transcriptional activity using scRNAseq on immune cells isolated from mouse brains over a time course of VEEV infection. We first confirmed infection dynamics in an established mouse model of VEEV. C3H mice were infected intranasally with 2×10^7^ PFU VEEV TC-83 and monitored for signs of disease over six days, encompassing both the early lymphoid phase and central nervous system phase of the typically biphasic disease observed in mice [17, 18]. Signs of disease including ruffled fur and hunched posture were observed in some mice by 4 days post infection (dpi) and by 6 dpi all mice exhibited these symptoms in addition to ataxia, a sign of neurological disease (**Fig. 1a**). Weights of infected mice began to decline by 5 dpi and were significantly different versus uninfected controls by 6 dpi (**Fig. 1b**). At various timepoints post infection, additional groups of mice were euthanized to measure brain viral load. High infectious viral titers in the brain were detected by 2 dpi and peaked by 4 dpi (**Fig. 1c**). Histological examination of infected brains was performed at 6 dpi. Hematoxylin and eosin-stained coronal sections of infected brains revealed increased cellularity, edema, and perivascular cuffing (**Fig. 1d, e**). Our results closely recapitulated previously characterized disease progression and pathological characteristics in this model of VEEV infection [18].

**Fig. 1:**
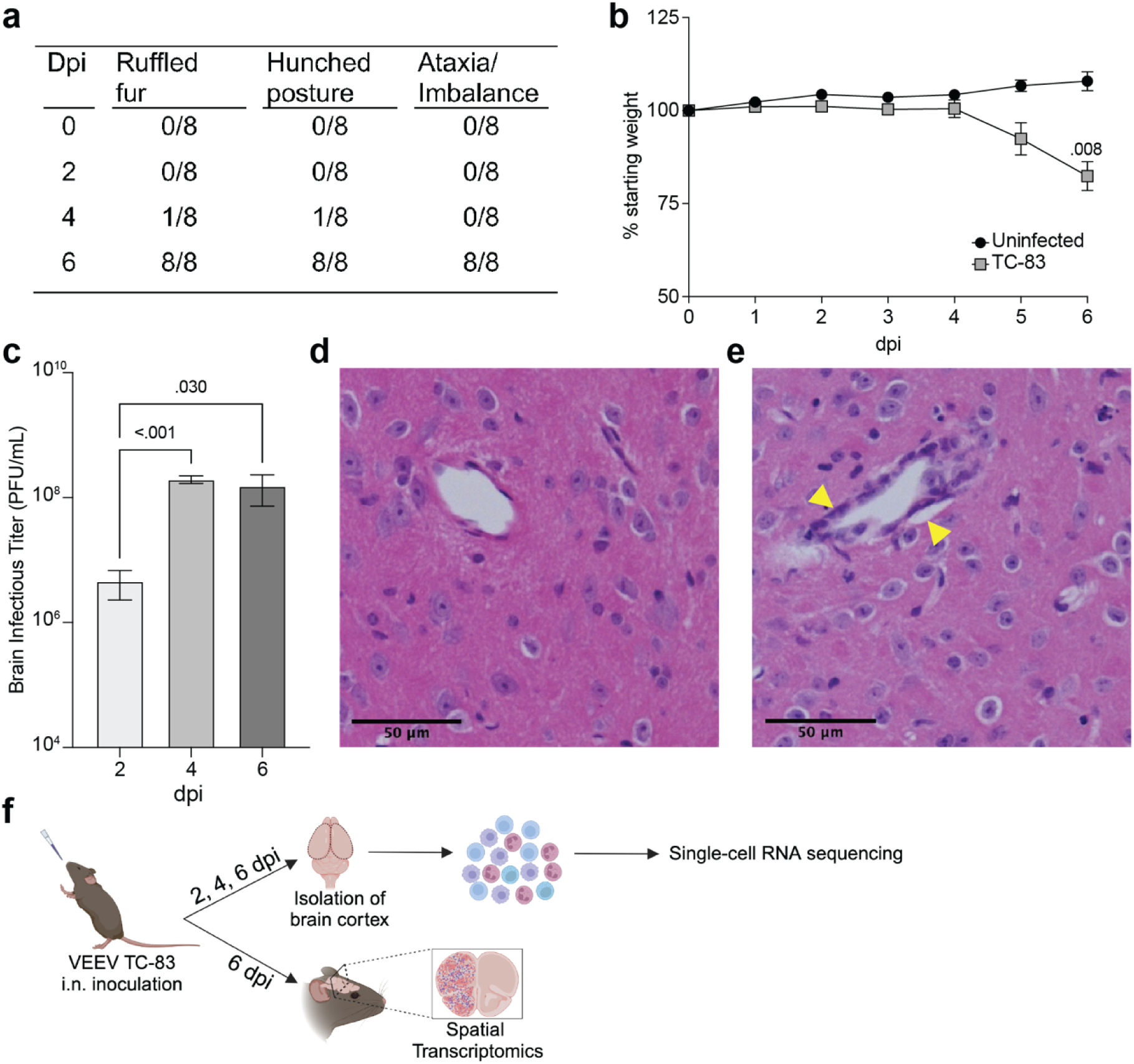
VEEV TC-83 Infection in C3H mice. 5–8-week-old C3H mice were infected intranasally with 2e7 PFU of VEEV TC-83 and symptoms (**a**) and weight loss (**b**) were monitored. Data are shown as mean ± standard error of mean (SEM) (*n*=4-12). Experiments were repeated at least twice. Two-tailed *p* values were calculated using 2-way ANOVA with Bonferroni’s correction for multiple comparisons. Brain viral titers were evaluated (**c**). Two-tailed *p* values were calculated using Kruskal-Wallis and Dunn’s multiple comparisons test. Signs of pathology in the brain were observed by H&E histological staining (**d**, **e**); yellow arrows denote vascular cuffing. (**f**) Schematic representation of experimental workflow for scRNAseq and spatial transcriptomic analysis of VEEV-infected brains. Following VEEV infection as described above, the cerebral cortex of 3 uninfected mice or 3 infected mice harvested at 2, 4, and 6 dpi were processed to yield a single cell suspension of mononuclear immune cells that were analyzed by scRNAseq. Alternatively, uninfected and infected cerebral cortices harvested at 6 dpi were fixed and processed for analysis by spatial transcriptomics. Created with BioRender.com.

C3H mice were then infected as described above and at 2, 4, and 6 dpi, brain cerebral hemispheres from infected mice and uninfected controls were harvested and processed to perform scRNAseq or spatial transcriptomic analyses (**Fig. 1f**). For scRNAseq, single-cell suspensions from 3 mice were pooled and mononuclear immune cells were isolated using Percoll gradient centrifugation, as previously described [19]. The following number of cells were profiled for each condition: Uninfected: 2,298, TC-83 2 dpi: 1,698, TC-83 4 dpi: 5,005, and TC-83 6 dpi: 6,488. Unsupervised clustering of the data resulted in 9 clusters that were each assigned to a putative cell-type identity based on their unique profile of differentially expressed genes (DEGs) encoding cell-type specific markers (**Fig. 2a, b**). Brains at each timepoint post-infection exhibited unique immune compositions as compared to the uninfected control (**Fig. 2c**). In uninfected brains, microglia were the predominant immune population detected, as expected. Microglia formed 3 subclusters over the course of infection that we termed resting microglia (MG), activated MG 1, and activated MG 2 (microglia 1, 2, and 3). Activated MG 1 and 2 are representative of a deviation from a homeostatic cell state upon infection. Resting MG compose 88% of total immune cells in uninfected mice and are the most enriched in microglia homeostatic markers *Cx3cr1* and *Tmem119*, which are downregulated in activated MG 1 and activated MG 2 (**Fig. 2b, Supplementary Fig. 1a**). At 2 dpi, activated MG 1 and 2 increased in abundance (16% and 3%, respectively) and exhibit enrichment in additional genes indicative of an activated state, including *Cd63*, *Cd72*, and *Mif* (**Fig. 2b, D**). Activated MG 1 continued to increase in abundance through 4 dpi, detected at a comparable level to resting MG, 16% and 17%, respectively, while activated MG 2 remained at approximately 3%. At 6 dpi, activated MG 1 decreased to 3% and activated MG 2 to less than 1%. Distinct antiviral response gene expression was observed across the microglia subclusters, with activated MG 1 exhibiting the highest expression of viral RNA sensors *Ifih1,* which encodes melanoma differentiation-associated protein 5 (MDA5) and *Ddx58*, which encodes retinoic acid-inducible gene I (RIG-I), as well as type I IFN gene *Ifnb1* (**Supplementary Fig. 1a**). Following the early shifts in microglial populations, we observed an immense infiltration of myeloid and lymphocyte populations that together comprise a larger proportion of sequenced cells at day 4 and 6 post infection than microglia (**Fig. 2c, d**). We additionally confirmed this pattern of myeloid and lymphocyte infiltration using flow cytometry (**Fig. 2e, f)**.

**Fig. 2:**
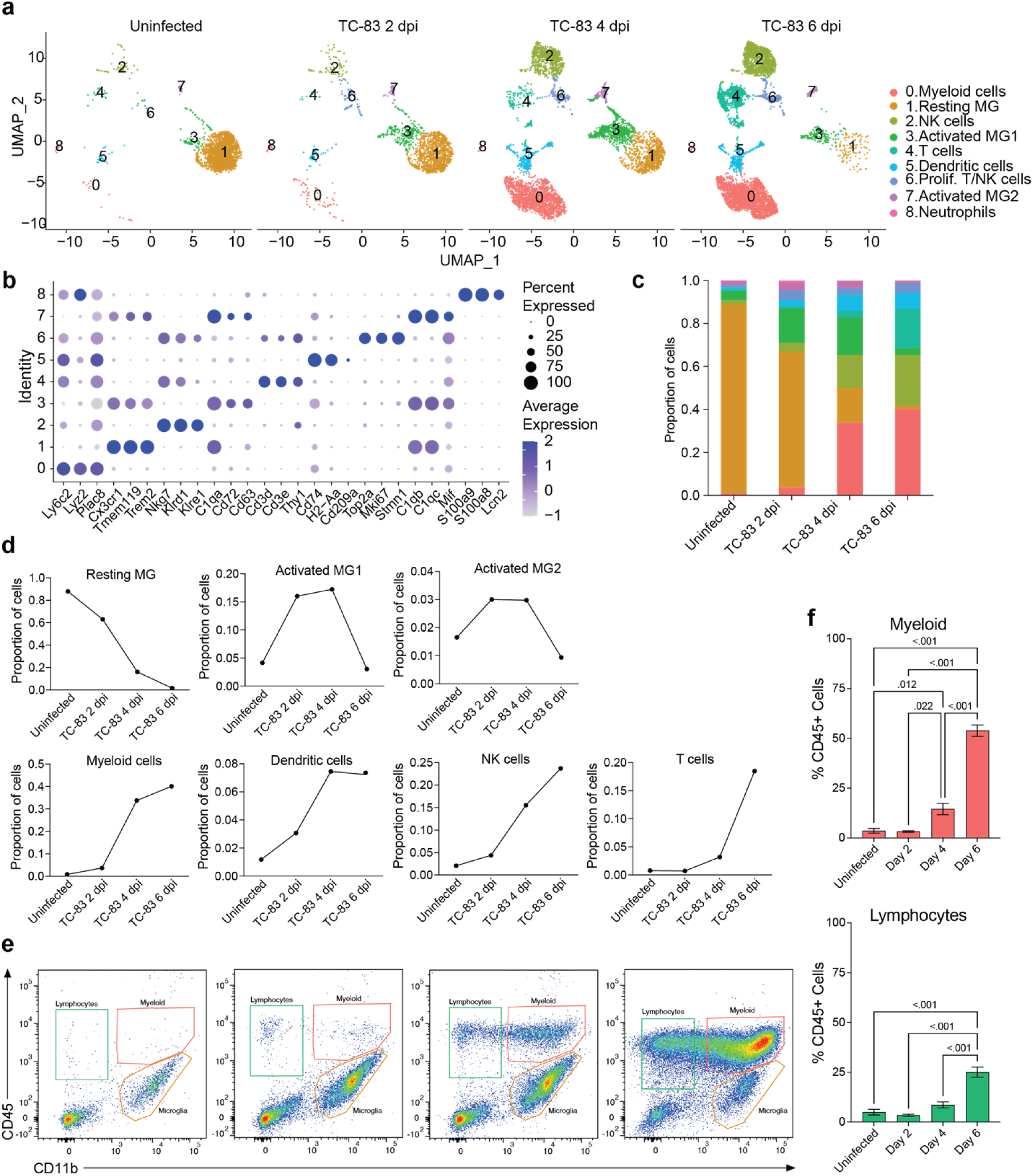
Temporal profiling of immune cells in the brain by scRNAseq in VEEV infection. (**a**) Uniform Manifold Approximation and Projection (UMAP) visualization of cell clusters identified in each experimental group. Cell clusters are color-coded and numbered. (**b**) Dot plot showing the expression of select cell-type markers. Dot size represents the fraction of cells of the indicated cluster expressing the markers and the intensity of color represents the average marker expression level in that cluster. (**c**) The relative proportion of cell types within brains of each experimental group. (**d**) Line graph representation of the change in proportion of prominent cell types over time in uninfected and infected brains. (**e**) Flow cytometric analysis of cell types in uninfected and infected brains. Representative CD11b vs CD45 plots are shown with gates identifying lymphocytes (CD11b^−^, CD45^hi^), myeloid cells (CD11b^hi^, CD45^hi^), and microglia (CD11b^hi^, CD45^low^) (**f**) Percentages of lymphocytes and myeloid cells. Data are shown as mean ± SEM (*n*= 4-7). Experiments were repeated at least twice. Two-tailed *p* values were calculated using one-way ANOVA with Tukey’s correction for multiple comparisons.

Between uninfected and 2 dpi, the most notable shifts were those in the microglia subclusters, as described earlier, as well as detection of cluster 0, identified as myeloid cells, cluster 2, identified as NK cells, and cluster 6, identified as proliferating NK and T cells. Between 2 and 4 dpi, there was a large increase in myeloid cells (4% to 34%) and NK cells (4% to 16%) and a modest increase in cluster 5, identified as dendritic cells (DC) (3% to 7%). We also identified a small neutrophil cluster which accounted for less than 0.5% of cells at 4 and 6 dpi. T cells accounted for ∼3% of the immune cells present in the brain at 4 dpi. Between 4 and 6 dpi, the myeloid and dendritic cell clusters remain relatively steady (40% and 7%, respectively) while there is an increase in NK cells (16% to 24%) and cluster 4, identified as T cells (3% to 18%) (**Fig. 2d, e**). Notably, it is between day 4 and 6 during which neurological symptoms and severe brain pathology present and weight loss and survival begin to decline.

Williams et al. recently investigated the dynamics of the host response to VEEV infection in the mouse brain using bulk RNA sequencing (RNAseq) and histological techniques [20]. We reanalyzed the bulk RNA-seq data obtained from three different regions of the brain: 1) the main olfactory bulb (MOB) where robust expression of viral protein was detected as early as 1 dpi, 2) piriform cortex (PIR), a region with robust viral protein expression by 2 dpi and 3) hippocampus (HIP), a region showing strong viral protein expression by 4 dpi [20]. This data showed that *Ifih1* and *Ddx58* were upregulated immediately after infection in all three regions, which was followed by upregulation of type I IFN genes including *Ifna2*, *Ifna4*, *Ifna5* and *Ifnb1* (**Supplementary Fig. 1b**). Interferon expression decreased over time in all three regions. Consistent with the decrease in resting microglia observed in our scRNAseq data, a dramatic decrease in the expression of microglia markers *Cx3cr1*, *Tmem119*, *P2ry12* and *Fcrls* was observed while several markers of infiltrating myeloid cells including *Ly6c2* and *Plac8* were upregulated (**Supplementary Fig. 1b**). In addition, a significant increase in the expression of NK and T cell markers over time was also observed in all three regions examined. In alignment with our data, an increase in NK cell markers was observed between 3 and 7 dpi, followed by a significant increase in T cell markers between 5 dpi and 7 dpi (**Supplementary Fig. 1b**). Consistent with the lack of B cells and few neutrophils observed in our scRNAseq data, the expression of B cell and neutrophil markers was extremely low, as shown in the MOB at 6 dpi (**Supplementary Fig. 1c**). We then reanalyzed the expression of interferon and markers of the key infiltrating populations in all eight parts of the brain isolated by Williams et al., (MOB, PIR, striatum (STR), motor cortex (MTX), HIP, sensory cortex (STX), thalamus (THA), and cerebral cortex (CBX). High expression of type I IFN genes was observed earliest in the MOB and PIR, at 3 dpi, followed by STR, MTX, HIP, STX, and THA across 5 and 6 dpi **(Supplementary Fig. 1d**). *Ly6c2 was* upregulated in the MOB from 3 through 7 dpi whereas it was upregulated from 5 to 7 dpi in the PIR, STR, MTX, HIP, STX, and THA. NK cell and T cell markers were upregulated highest from 5 to 7 dpi in the MOB, and PIR, and at 7 dpi in the MTX, HIP, STX, and THA. Expression of these markers was least changed in the CBX across all timepoints. The early expression of viral sensing genes and type I IFN genes in the bulk RNAseq data set preceding high expression of markers of infiltrating cells coincides with our scRNAseq data that implicates microglia as a key early responder to VEEV infection and early producer of interferon. Both data sets then indicate a sequential recruitment of myeloid cells followed by lymphocytes at later timepoints. The bulk RNAseq data set demonstrates that in addition to temporal differences, there are unique spatial signatures of gene expression that can potentially help define the progression of the immune response through specific regions of the brain over time and the extent of infiltration in each of these regions.

### Microglial and myeloid cell subtypes elicit heterogeneous antiviral responses following VEEV infection

To further examine the transcriptional activity of microglia and infiltrating myeloid populations, all cells from clusters 0, 1, 3, 5, 7, and 8 (**Fig. 2a**) were extracted and re-clustered using an unsupervised clustering approach. This included the following number of cells for each condition: Uninfected: 2,211, TC-83 2 dpi: 1,527, TC-83 4 dpi: 3,835, and TC-83 6 dpi: 3,398. Analysis revealed 10 unique cell clusters (**Fig. 3a-c**). Microglial subpopulations (cluster 0; resting MG, cluster 3; activated MG 1, and cluster 5; activated MG 2) clustered apart from various infiltrating myeloid populations and exhibited enrichment for *Cx3cr1*, *Trem2*, and *Tmem119*, as well as complement component genes *C1qa-c*. Cluster 1 and 2 highly expressed monocyte/macrophage markers *Ly6c2* and *Plac8*, while lacking *Tmem119* and were, therefore, annotated as ‘monocyte/macrophage (Mono/Mac) 1’ and ‘Mono/Mac 2,’ respectively (**Fig. 3c**, **Supplementary Fig. 2a**) [21, 22]. Clusters 4, 7, and 8 were identified as Cd209a^+^, plasmacytoid, and Ccr7^+^ DCs, respectively, based on enrichment of specific genes such as *Cd209a* and class II MHC genes *Cd74*, *H2-Aa* and *H2-Ab1* (cluster 4), *Klk1* and *Mctp2* (cluster 7) and *Ccr7* (cluster 8) (**Fig. 3c, Supplementary 2**) [23]. Cluster 9 expressed neutrophil markers *S100a8* and *S100a9* (**Fig. 3c**).

**Fig. 3:**
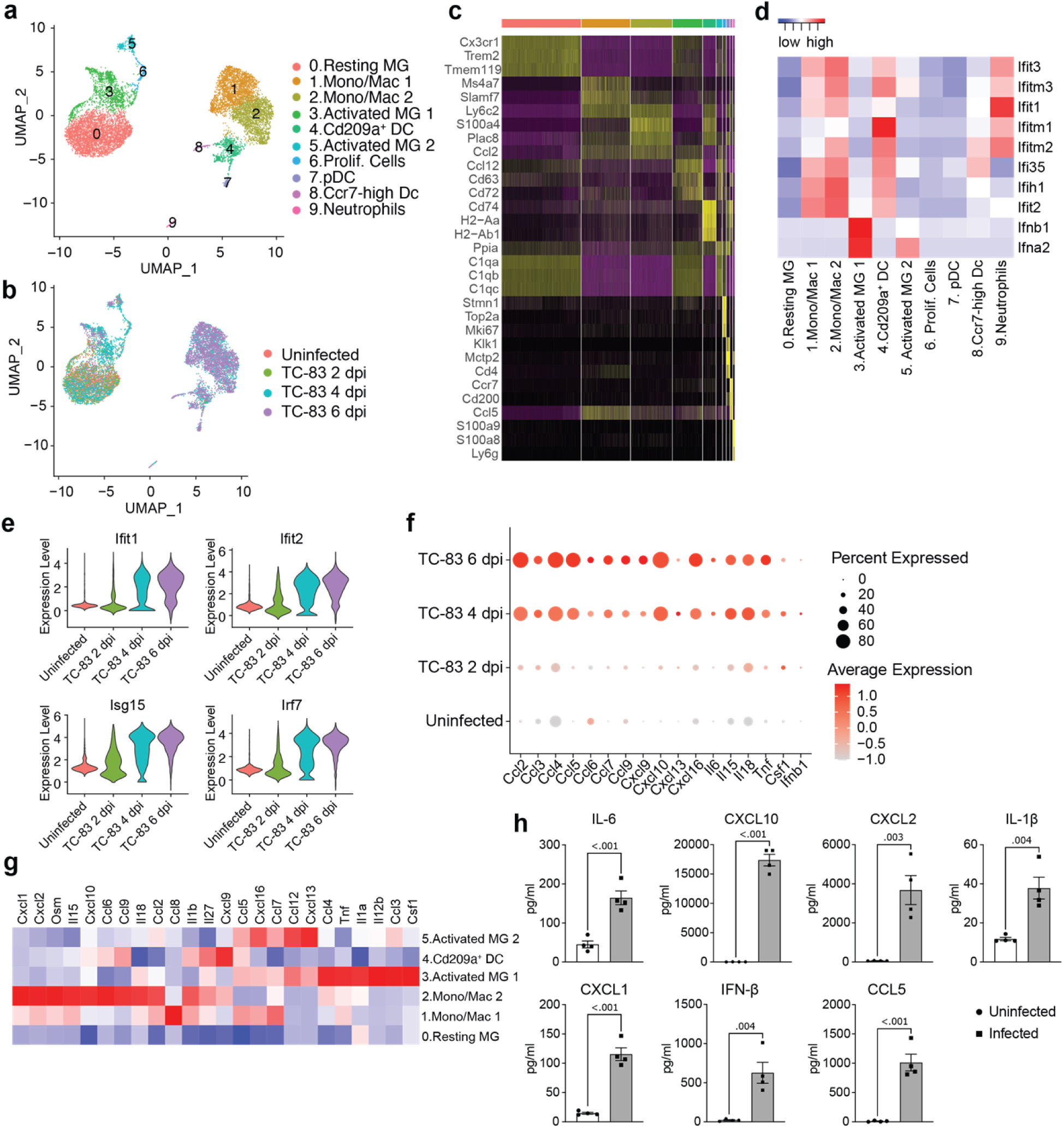
Analysis of myeloid populations in the brain in by scRNAseq in VEEV infection. UMAP visualization of myeloid sub-clusters colored by identification (**a**) or experimental group (**b**). (**c**) Heat map showing expression level of selected cell type markers in each cluster. (**d**) Heat map of IFN ⍺/β response genes. (**e**) Violin plots showing the expression of select ISGs across experimental groups. (**f**) Dot plot showing expression level of select inflammatory signaling genes across experimental groups. Dot size represents the fraction of cells in the indicated group expressing the indicated gene and the intensity of color represents the average marker expression level in that group. (**g**) Heat map showing expression level of select inflammatory signaling genes across key myeloid populations. (**h**) Protein levels of select immune mediators measured using BioLegend LEGENDplex multiplex immunoassay. Data are shown as mean ± SEM (*n*=4). Two-tailed *p* values were calculated using unpaired Student’s t test.

To understand the function of the myeloid clusters, we analyzed expression of key antiviral response genes. Type I IFN genes *Ifnb1* and *Ifna2* were primarily expressed by microglia, specifically activated MG 1 (**Fig. 3d)**. Elevated levels of interferon stimulated genes were observed across other myeloid subclusters, mainly Mono/Mac 1 and 2 and Cd209a^+^ DCs as early as 2 dpi, but more notably at 4 dpi and through 6 dpi (**Fig. 3d, e**). Activated MG 1 were also enriched for inflammatory cytokines *Ccl3* and *Ccl4*, *Csf1*, a key regulator of macrophage differentiation, tumor necrosis factor alpha (*Tnf*), and *Il12b*, a cytokine that acts on T and natural killer cells (**Fig. 3g**). *Ifnb1* and *Csf1* elevation was observed by 2 dpi and through 6 dpi, in a small (< 20) percent of total cells (**Fig. 3f)**, while the other cytokines were observed highly expressed by greater percentages of total cells and beginning at 4 dpi, through 6 dpi. Activated MG 2 exhibited elevated *Ifna2* expression, though lower than activated MG 1 and not until 6 dpi (**Fig. 3d, Supplementary Fig. 1a**), and were enriched for *Ccl5*, *Ccl7*, *Ccl12*, *Cxcl16*, and *Cxcl13* (**Fig. 3g**). Mono/Mac 1 shared overlap with activated MG 2 with enrichment for *Ccl5*, *Ccl7*, and *Cxcl16*, and additionally expressed high levels of *Ccl8*. Mono/Mac 2 expressed high levels of chemokines including *Ccl2*, *Ccl6*, *Ccl9*, *Cxcl1*, *Cxcl2* and *Cxcl10*, as well as *Il15* and *Il18*, two interleukins implicated in activation of T/NK cells (**Fig. 3g**) [24–26]. Mono/Mac 1 also exhibited elevated expression of *Il15* and *Il18*, but to a lesser extent. Notably, Cd209a^+^ DCs had high expression of *Cxcl9,* a prominent mediator of lymphocyte infiltration (**Fig. 3g**). We validated our transcriptomic data by measuring protein levels of a subset of cytokines and chemokines included in our analysis in brain homogenate isolated from infected mice. Consistent with the observed increase in gene expression, IL-1β, IL-6, CXCL1, CXCL2, CXCL10, CCL5 and IFN-β were upregulated at the protein level in infected brains (**Fig. 3h**).

To investigate the relationship between myeloid subclusters, we performed trajectory analysis, ordering single cells along pseudotime to reconstruct potential differentiation trajectories (**Supplementary Fig. 2b**). Mono/Mac and microglia subpopulations clustered distinctly through each timepoint. Microglia clusters exhibited a shift from resting to activated subpopulations. By 4 dpi, activated MG 1 partially occupy a shared branch with Mono/Mac clusters. Meanwhile, activated MG 2 and proliferating cells occupy distinct branches. Interestingly, only a small portion of activated MG 1 highly express *Ifnb1* at each timepoint (**Supplementary Fig. 2c**).

### Infiltrating myeloid cells occupy specific spatial niches in the brain

Having established that microglia subclusters and infiltrating myeloid cells display distinct inflammatory profiles in response to VEEV infection, we next asked whether the spatial distribution of these immune cell subtypes also differ in the brain parenchyma using spatial transcriptomic analysis performed at 6 dpi. Unsupervised clustering of the spatial transcriptomic data from uninfected and infected brains resulted in eleven clusters. The majority of these clusters corresponded to specific anatomical regions in the brain (**Fig. 4a-c**). Cluster 2 showed enrichment for myeloid markers such as *Ly6c2* and *Plac8* and was localized at the periphery of the cortex and within the hippocampal region (**Fig. 4c-f**). The uninfected brain had significantly fewer cells from cluster 2 compared to the infected brain at 6 dpi (**Fig. 4e, f**). Examination of spatial expression patterns of *Ly6c2* and *Plac8* revealed a distinct localization of myeloid cells expressing these genes also in the periphery of the cortex and within the hippocampal region of infected brains (**Fig. 4g**). Localized Ly6c was confirmed in these regions via immunohistochemical analysis (**Fig. 4i**). Minimal Ly6c expression was noted in the uninfected brain and robust signal was detected in the infected brain concentrated around the hippocampal formation (**Fig. 4i**). When compared to the localization patterns of *Prox1*, which appears enriched at the hippocampal formation, and *Nrgn1*, within the cortex, the expression of the myeloid-specific markers appears to distinctly localize at a layer in between the *Prox1*- and *Nrgn1*-high regions, within the hippocampal region (**Supplementary Fig. 3d**, **Fig. 4g**). Our re-analysis of the bulk RNAseq data set generated by Williams *et al*. also indicated greater immune infiltration, including myeloid populations, within the hippocampus compared to the cortex (**Supplementary Fig. 1d**). Expression of several upregulated chemokines in the infected brain including *Ccl2*, *Ccl4*, *Ccl5*, *Ccl7* also localized to regions enriched in myeloid cells whereas *Tnf* and interleukins including *Il6*, *Il12b*, *Il15* and *Il18* were detected throughout the brain (**Fig. 4c, Supplementary Fig. 3a, b**). Interestingly, *Cxcl10* had a more robust expression throughout regions of the infected brain other than the *Ly6c* enriched regions, suggesting that *Cxcl10* may be primarily expressed by cells other than the localized myeloid subpopulation (**Supplementary Fig. 3c**). Consistent with the scRNAseq data, there was a significant reduction in cells expressing markers of homeostatic microglia such as *Cx3cr1* and *Tmem119* and an increase in cells expressing *Cd72*, a gene enriched in activated microglia, in infected brains (**Fig. 4h**). These data indicate that by 6 dpi, there is widespread activation of microglia and regional specificity in infiltrating myeloid subpopulations.

**Fig. 4:**
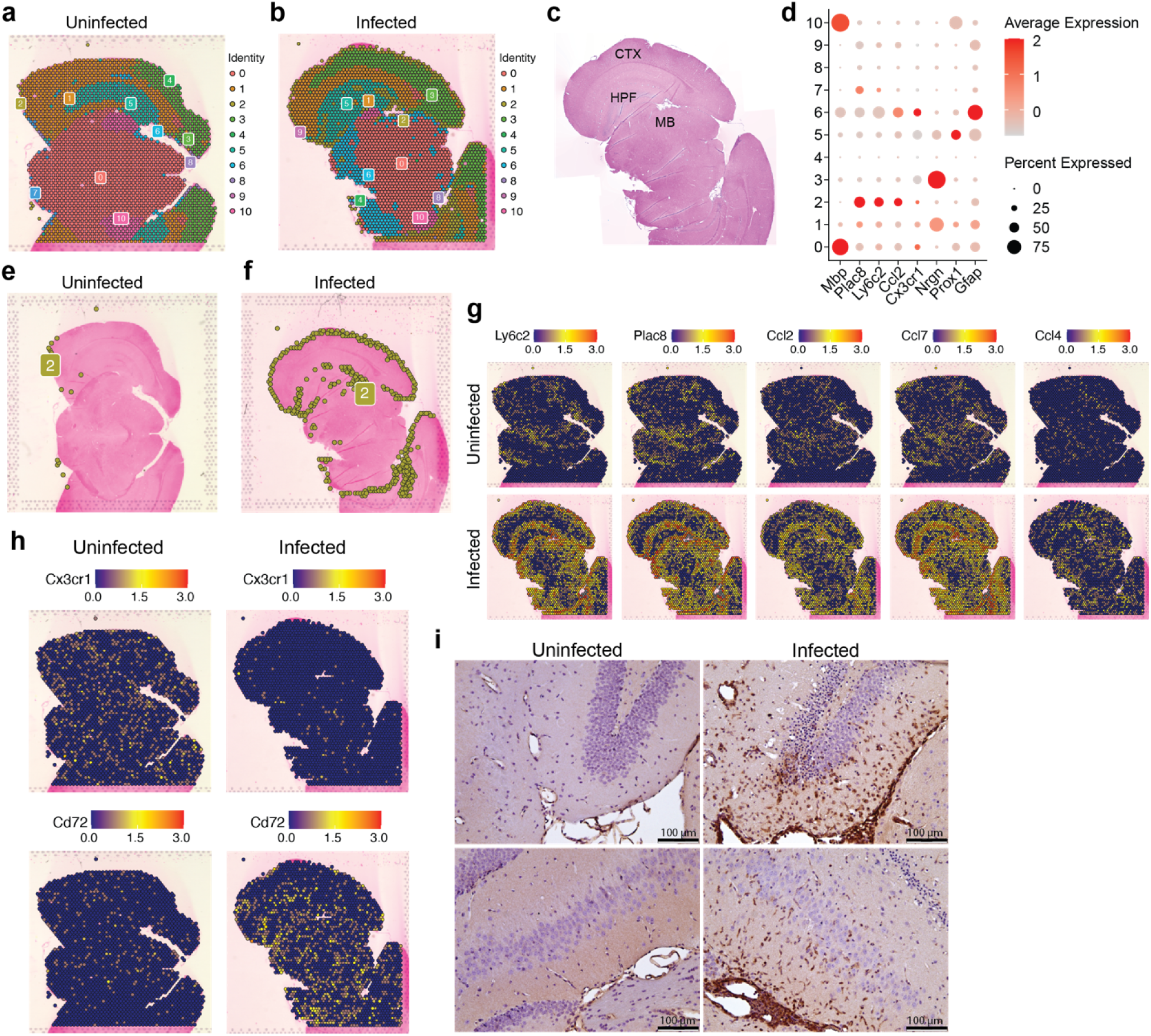
Spatial transcriptomic profiling of myeloid populations in the brain during VEEV infection. 5-μm coronal sections through the hippocampus were derived from uninfected mice or infected mice at 6 dpi. Sections were used for either ST (**a**-**h**) or immunostaining (**i**). Distribution of transcriptionally unique populations detected in uninfected (**a**) or infected (**b**) brains. Populations are numbered and color-coded. (**c**) Annotation of major brain regions. MB= midbrain, HPF= hippocampal formation, CTX= cortex. (**d**) Dot plot showing expression levels of identifying markers of cell types across the brain. Dot size represents the fraction of cells in the indicated cluster expressing the markers and the intensity of color represents the average marker expression level in that cluster. Distribution of cluster 2 is shown in uninfected (**e**) or infected (**f**) brains. Distribution of select myeloid gene and cytokine expression (**g**) and microglial gene expression (**h**). (**i**) Representative images of Ly6C staining in the hippocampus (magnification = 20).

### Recruited NK and CD8 T cells showed a widespread distribution in the brain parenchyma and elevated expression of cytotoxic proteins

To gain a more comprehensive understanding of lymphocyte responses to infection, all cells from lymphocyte clusters (clusters 2, 4 and 6; **Fig. 2b**) were extracted and reanalyzed. This included the following number of cells for each condition: Uninfected: 66, TC-83 2 dpi: 89, TC-83 4 dpi: 987, and TC-83 6 dpi: 2783. Cells were primarily identified as one of two subclusters of NK cells, referred to as NK cells 1 and 2, CD8^+^ T cells, CD4^+^ T cells, or smaller detected clusters of γδT cells and cells expressing markers of proliferation (**Fig. 5a**). At both 4 and 6 dpi, NK cell clusters composed a greater proportion of total lymphocytes than CD8^+^ and CD4^+^ T cells, which exhibited a steep increase at 6 dpi (**Fig. 5b, c**). High expression of genes encoding cytotoxic proteins, including perforin, granzyme B, and FasL, were observed at both 4 and 6 dpi (**Fig. 5d**). Perforin and granzyme genes (*Prf1* and *Gzmb*, respectively) were expressed in both NK subclusters and CD8^+^ T cells at both timepoints, with NK cells exhibiting higher expression (**Fig. 5e**). While expression in NK cells was comparable for both timepoints, CD8^+^ T cells exhibited higher expression of *Prf1* and *Gzmb* at 6 dpi. Additionally, *Ifng* expression was detected in both NK sub clusters and CD8^+^ T cells, each exhibiting higher expression at 4 dpi. Production of perforin at 6 dpi was also observed by flow cytometry and IFN-γ by protein analysis (**Fig. 5g-j**). Perforin protein was detected in NK cells and CD8^+^ T cells, but not CD4^+^ T cells (**Fig. 5i**). Further, perforin protein was significantly higher in the brain than spleen in both cell types, suggesting unique cytotoxic effector expression in the brain versus systemically. An increase in the expression of NK cell inhibitory genes such as *Klrc1* and *Klrd1* was observed in both NK and CD8^+^ T cells at 6 dpi (**Fig. 5e**). This together with the decreased *Ifng* expression at 6 dpi may indicate NK cell exhaustion around the later timepoint. Spatial transcriptomic analysis at 6 dpi revealed detection of NK and T cell associated genes throughout the brain with no observable specific localization patterns, though, *Ifng* expression sometimes formed concentrated clusters (**Fig. 5f**). These results indicate a robust and ubiquitous upregulation of cytotoxic effectors throughout the brain in response to VEEV infection that coincide with timepoints of severe neurological dysfunction.

**Fig. 5:**
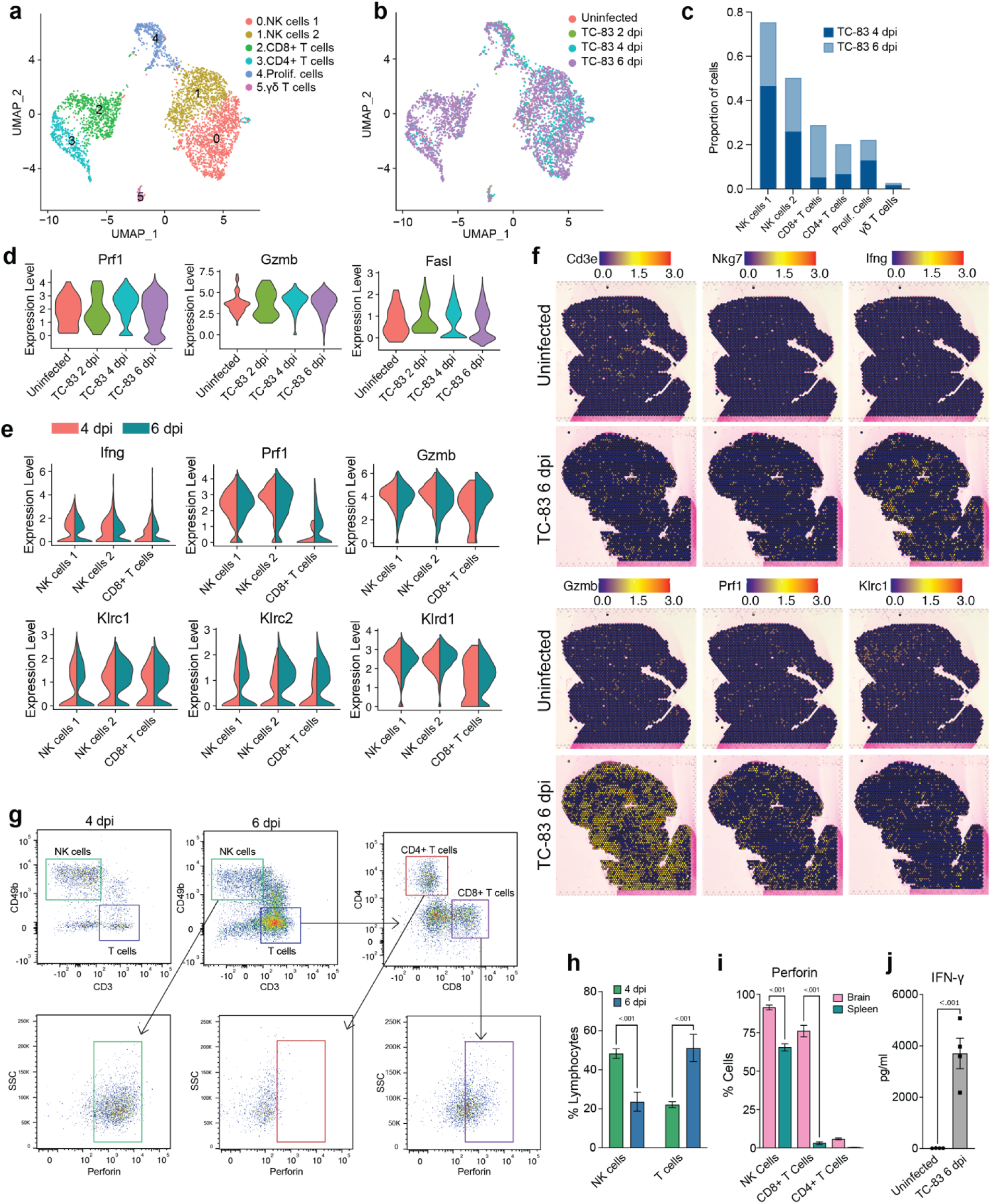
Analysis of lymphocyte populations in the brain by scRNAseq and spatial transcriptomics. UMAP visualization of lymphocyte clusters colored by identity (**a**) or experimental group (**b**). (**c**) Proportions of lymphocyte populations. (**d**) Violin plots showing expression of cytotoxic genes across experimental groups. (**e**) Violin plots showing expression of select NK and T cells genes within the indicated subpopulations. Colors indicate timepoint post infection. (**f**) Spatial distribution of select NK and T cell genes in uninfected or infected brains at 6 dpi. (**g**) Representative flow cytometry plots showing perforin expression in NK and T cell populations in the brain at 6 dpi. Quantification of NK and T cells in the brain is shown in (**h**) and perforin in the brain versus spleen in (**i**). (**j**) Protein analysis of IFN-γ in uninfected or infected brains at 6 dpi. Data in **h**-**j** are shown as mean ± standard error of mean (SEM) (*n*=4-7). Two-tailed *p* values were calculated using 2-way ANOVA with Bonferroni’s correction for multiple comparisons (**h, i**) or unpaired Student’s t test (**j**).

### Comparative analysis of infected brain and peripheral blood mononuclear cells reveals distinct immune landscapes

To compare the immune landscape in the brain to systemic immune changes induced by VEEV infection, we profiled peripheral blood mononuclear cells (PBMCs) from infected and uninfected mice at 6 dpi using scRNAseq. Our analysis identified twelve immune cell clusters including NK cells, T cells, B cells, mono/macs, neutrophils, DCs and plasma cells (**Fig. 6a-c**). Consistent with our findings in the brain, the proportion of NK cells dramatically increased in the blood after infection (**Fig. 6d**). Mono/Macs also expand in both the infected blood and brain (**Fig. 3c, d and 6d**). In contrast to what we observed in the brain, T cells 1, which encompassed CD4^+^ T cells and regulatory T cells, decreased and the proportion of CD8^+^ T cells remained similar between infected and uninfected blood (**Fig. 6d**). Neutrophils, which do not infiltrate the brain, dramatically expanded in the blood after infection and upregulated antiviral response genes, including *Ddx58*, *Ifih1*, *Ifit1*, *Ifit2* and *Isg15* (**Fig. 6e**). Blood Mono/Macs and DCs also activated a large number of antiviral response genes in response to infection (**Fig. 6e**).

**Fig. 6:**
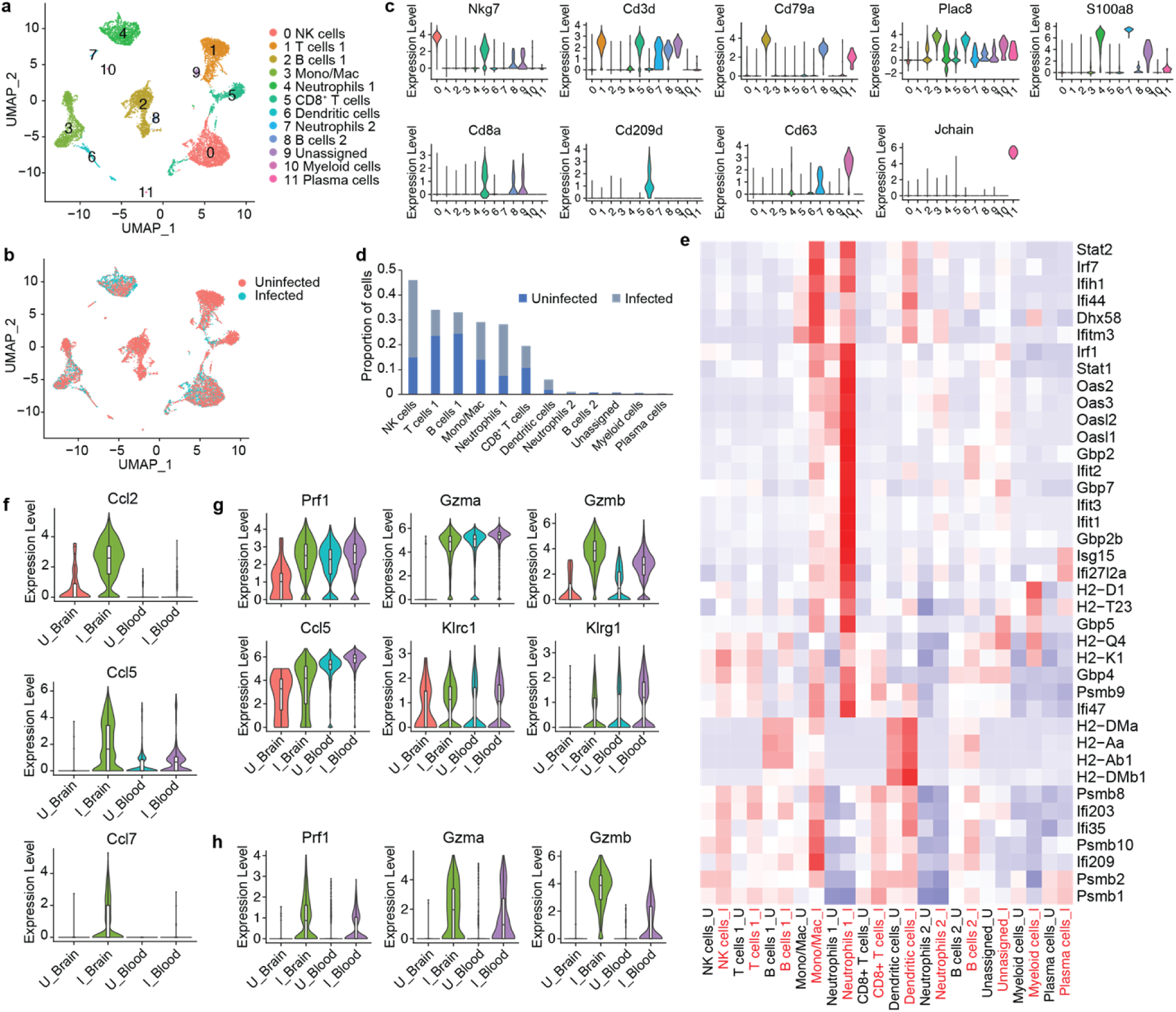
Comparative scRNAseq analysis immune cells in the brain and PBMCs during VEEV infection. UMAP visualization cell clusters in the blood at 6 dpi colored by identity (**a**) or treatment (**b**). **c** Expression of identifying markers across clusters. **d** Relative proportions of populations. **e** Heat map showing expression of select antiviral response genes across cell populations in uninfected versus infected blood. Comparison of the expression of select genes in mono/mac (**f**), NK cell (**g**) and CD8^+^ T cell (**h**) populations in the brain versus blood.

Next, we compared infection-induced transcriptional changes in PBMCs to immune cells from the brain **(Supplementary Fig. 4 a, b**). Similar to the brain, blood Mono/Macs from infected mice activated *Plac8*, *Ly6c2* and cathepsin family proteases (**Supplementary Fig. 4c**). We found that cytokines such as *Ccl2, Ccl5* and *Ccl7* had markedly lower expression in infected blood Mono/Macs compared to brain Mono/Macs while *Ccr2*, a receptor involved in myeloid recruitment to the CNS, had an elevated expression in blood Mono/Macs (**Fig. 6f, Supplementary Fig. 4c**) [27]. Peripheral blood NK cells from infected mice expressed perforin, granzymes, cytokines and inhibitory receptors at a level comparable to NK cells from infected brains, with slightly higher *Gzmb* expression in the infected brain (**Fig. 6g**). Interestingly, uninfected blood NK cells also had robust expression of *Prf1* and *Gzma* but, not *Gzmb*. The expression levels of perforin and granzymes were also upregulated in CD8^+^ T cells after infection (**Fig. 6h**). Unlike uninfected NK cells, uninfected blood CD8^+^ T cells did not express *Prf1* or *Gzma,* suggesting that cytotoxic gene expression in CD8^+^ T cells was infection-induced (**Fig. 6h**). Overall, these findings emphasize a tissue-specific upregulation of cytotoxic effectors as a potential key component of the immune landscape in the brain in response to VEEV infections.

## Discussion

Existing and emerging encephalitic viral infections pose a growing medical challenge. Viral infection of the brain and subsequent inflammatory immune responses can cause long lasting or permanent damage leading to serious clinical outcomes and death. Lethal encephalitis that ensues following VEEV infection is attributed to the host immune response to viral replication in brain tissue, but it remains unclear which specific components of the immune response instigate disease. Thus, understanding the processes involved in the recruitment, differentiation, and function of immune cells in the brain during infection is of high importance to aid the development of novel therapeutics that target immunopathological responses. Here we temporally dissected the heterogeneity of the immune response in the brain over a time course of infection using scRNAseq. We then employed spatial transcriptomic analysis of the brain during severe disease to investigate the spatial organization of distinct populations within the brain.

We observed a dynamic response to VEEV in myeloid cells over the course of infection. The emergence of two subpopulations of microglia was observed by 2 dpi that we termed activated microglia (MG) 1 and 2. Trajectory analysis revealed their simultaneous expansion, suggesting they are uniquely activated subpopulations. Notably, activated MG 1 was the primary cell type expressing genes encoding RIG-I (*Ddx58)* and MDA5 (*Ifih1)*, PRRs important in alphavirus sensing, along with type I IFN genes *Ifnb1* and *Ifna2* throughout infection. Activated MG 2 did not exhibit elevated *Ddx58* or *Ifih1* but showed elevated *Ifna2* expression at 6 dpi. Interestingly, within activated MG 1, only a small portion of the population expressed *Ifnb1* at each timepoint. While VEEV primarily infects neurons and astrocytes, it can also infect microglia to a lesser frequency [10]. It is possible that the unique expression profiles among microglia may represent infected cells versus bystander cells assuming an antiviral state. Alternatively, concurrent molecular pathways could be contributing to limiting *Ifnb1* expression. Further, a lack of upregulation of additional interferon-stimulated genes (ISGs) in the activated microglia subsets suggests their activation may be orchestrated by other cell types that are directly sensing virus.

The early and sustained activation of microglia during VEEV infection likely plays a role in the recruitment of other inflammatory cell types. Activated MG 1 and 2 share expression of cytokines expressed by disease-associated microglia (DAM) with important roles in the recruitment of inflammatory monocytes [28–30]. Ly6C^hi^ monocytes have been shown to play both protective and pathogenic roles upon migration to the brain [31–36]. In our data set, greater than 30% and 40% of immune cells in the brain are infiltrating myeloid cells by 4 and 6 dpi, respectively, consisting mostly of two *Ly6c2* expressing subpopulations (Mono/Mac 1 and 2) that display high induction of numerous ISGs, cytokines, and chemokines. Trajectory analysis indicated Mono/Mac 1 and 2 cluster separately throughout each timepoint, likely ruling out either population being a precursor state to the other. Notably, this large myeloid infiltration coincides with peak viral replication and the appearance of clinical signs and future studies should investigate the mechanistic role of these cells in VEEV infection.

Other notable myeloid cell populations that responded to infection include neutrophils and DCs. Both cell populations expanded in the blood and expressed high levels of antiviral genes in response to infection, yet only DCs were detected at appreciable levels in the brain. The lack of neutrophils in the brain is notable given their dramatic expansion in the blood and their significant infiltration of the brain during other neurotropic infections, including with other neurotropic arboviruses [37]. The expansion of neutrophils in the blood and their high expression of antiviral response genes may indicate a role in systemic viral clearance.

Lymphocyte populations exhibited diverse responses upon infection, with subset populations demonstrating brain-specific cytotoxic signatures. NK cells expanded considerably in the brains and blood of infected animals, and T cells exhibited a sharp increase in the brain at 6 dpi. Conversely, B cells decreased proportionally in the blood upon infection and did not infiltrate the brain. Both NK cells and CD8^+^ T cells from infected animals expressed elevated levels of cytotoxic genes, with higher expression observed within brain-infiltrating cells. While blood NK cells exhibited similar levels of *Prf1* and *Gzma* expression as in the brain, brain NK cells exhibited higher *Gzmb.* Notably, *Gzmb* exhibits a lower baseline expression in blood NK cells than *Gzma*, therefore the high level of *Gzmb* observed in the brain following infection represents a dramatic shift from baseline expression. Furthermore, striking differences in perforin expression in NK and CD8^+^ T cells were observed between cells isolated from brains versus spleens via flow cytometry, with brain populations of both cell types exhibiting significantly higher expression. Therefore, the highly cytotoxic state of brain infiltrating NK and CD8^+^ T cells is a specific quality and may be significantly contributing to immunopathology. Notably, previous work has implicated NK cells in the pathogenesis of VEEV in mouse models and the cytopathic effects of NK and T cells have also been implicated in neuroinflammation and blood brain barrier (BBB) disruption in non-infectious contexts [16, 38, 39]. The abundantly present cytotoxic lymphocytes are potentially key effectors of damage in the brain and, therefore, appealing therapeutic targets via blockade of their recruitment or function.

A key feature of our data is the unique spatial localization of infiltrating Mono/Mac cells within the brain on 6 dpi. A *Ly6c2-*expressing myeloid cluster was found to occupy a distinct regional pattern that lined the cortex perimeter and a region surrounding the hippocampal formation. In contrast, T and NK cells exhibited no clear localization pattern and were distributed throughout the brain. These data suggest fundamental differences in the mechanisms governing myeloid and lymphocyte infiltration into the brain. NK and T cells may require breakdown of the BBB in order to infiltrate in significant numbers, reflected in their widespread distribution throughout the brain, whereas myeloid cells may traverse alternate barriers (choroid plexus, glia limitans) at earlier time points. The mechanisms that control myeloid cell extravasation into the brain parenchyma across any barrier are under described [40], therefore additional research into entry mechanisms during viral infection is critical. The relationship between localized viral replication within the brain and the proximity to clustered immune populations also warrants further investigation. Notably, concentrated infected/apoptotic neurons have been observed in the hippocampus in VEEV-infected mice [41]. In turn, the *Lyc62-* expressing cluster likely contributes to further inflammation, as these cells exhibited high expression of chemokines, including *Ccl2*, *Ccl4*, and *Ccl7*. The role of each cell population and features such as their localization, expression profiles, and temporal infiltration during VEEV pathogenesis are of keen interest for follow on studies.

In summary, this study provides a comprehensive profiling of transcriptional activity of immune cells in the brain during viral encephalitis. We define the heterogeneity of the immune response in the brain at three timepoints post-VEEV infection, tracing the activation of resident cells, infiltration of cells from the periphery, and the emergence of various subpopulations. We compared the brain gene expression data to that of PBMCs to underscore brain-specific changes. Spatial transcriptomics identified localization of an infiltrating myeloid population to discrete regions of the brain, and furthermore highlighted distinct localization patterns of myeloid cells and lymphocytes. Future studies should utilize the data provided here to further define the immunopathogenic components of the response to VEEV versus those that are protective to inform the development of vaccine and therapeutic candidates.

## Materials and Methods

### Cells and virus

Vero E6 cells were obtained from the American Type Culture Collection (ATCC) and maintained in Dulbecco’s modified Eagle’s medium (DMEM, Thermo Fisher) supplemented with 10% fetal bovine serum (FBS, ATCC) supplemented with 100 units/mL penicillin and 100 μg/mL streptomycin (Thermo Fisher) at 37 °C in 5% CO_2_. Venezuelan equine encephalitis virus (VEEV) strain TC-83 (NR-63) was obtained from the NIH Biodefense and Emerging Infections Research Resources Repository, NIAID, NIH. VEEV stocks were propagated in Vero E6 cells and harvested via clarification of cell culture supernatant by centrifugation. Titers of viral stocks were determined by standard plaque assay consisting of a methyl crystalline cellulose overlay and crystal violet staining [42].

### Mouse infections

All animal work was approved by the Lawrence Livermore National Laboratory Institutional Animal Care and Use Committee under protocol #310. All animals were housed in an Association for Assessment and Accreditation of Laboratory Animal Care (AAALAC)-accredited facility. C3H/HeN mice (strain ID 025) were obtained from Charles River. 5–10-week-old female mice were used in all experiments. Groups of mice were inoculated intranasally with 2×10^7^ PFU TC-83 while under anesthesia (4-5% isoflurane in 100% oxygen). Mice were monitored daily for signs of morbidity and animals were humanely euthanized upon signs of severe disease by CO_2_ asphyxiation. For tissue harvest, animals were anesthetized under isoflurane and the whole animal was perfused with 20 mL sterile PBS containing 50,000 U/L sodium heparin *via* the left ventricle.

### Brain tissue isolation and preparation

Brains for flow cytometric, scRNAseq, cytokine, and viral titer analysis were isolated from infected mice on days 2, 4, or 6 post-infection. Preparation of brain tissue was performed as previously described [19]. Following euthanasia and perfusion, brains were removed and placed in 1 mL digestion buffer (PBS pH 7.4 (Thermo Fisher) + collagenase (Worthington) + DNase I (Roche) to a final concentration of 3 mg/mL and 0.5 mg/mL, respectively) on ice in a 1.5 mL tube. Brains were finely diced into 1-2 mm^3^ pieces with scissors and tissue was digested at 37 °C for 30 min in a total of 5 mL digestion buffer. A cell suspension was generated by gentle pipetting followed by passage through a 70 μm cell strainer. The cell strainer was rinsed with PBS supplemented with 5% FBS to a total volume of 20 mL. Aliquots of this suspension were stored at −80 °C for cytokine and viral titer analysis. The remaining suspension was subjected to Percoll gradient centrifugation to purify mononuclear immune cells for flow cytometric analysis or RNA sequencing as previously described [19].

### Single-cell RNA sequencing of brain samples and data analysis

Single-cell suspensions of mononuclear immune cells from uninfected mouse brains and infected brains were prepared as described above. Cells were counted on a Countess II automated cell counter prior to single-cell sequencing preparation using Chromium Single-cell 3ʹ GEM, Library & Gel Bead Kit v3 (10x Genomics Cat # 1000075) on a 10× Genomics Chromium Controller following manufacturers’ protocol. Subsequently, libraries were sequenced on Illumina NextSeq 2000. The Cell Ranger Single-Cell Software Suite (10x Genomics) was then used to perform sample demultiplexing, barcode processing, and single-cell gene counting. After demultiplexing the sequencing files with “cellranger mkfastq”, “cellranger count” was used to perform alignment to mouse reference transcriptome (mm10), barcode processing and gene counting. Further analysis was performed using Seurat [43]. First, cells with fewer than 500 detected genes per cell or mitochondrial content greater than 10% and genes that were expressed by fewer than 5 cells were filtered out. Potential doublets and CD45^−^ cells were also removed. After pre-processing, we performed data normalization, scaling, and identified 2000 most variable features. Then, anchors for data integration were identified using the ‘FindIntegrationAnchors()’ function. Next, these anchors were passed to the ‘IntegrateData()’ function and a new integrated matrix with all four datasets was generated. After data integration, data was scaled, and the dimensionality of the data was reduced by principal component analysis (PCA). Subsequently, cells were grouped into an optimal number of clusters for de novo cell type discovery using Seurat’s ‘FindNeighbors()’ and ‘FindClusters()’ functions. A non-linear dimensional reduction was then performed via uniform manifold approximation and projection (UMAP) and various cell clusters were identified and visualized. Genes differentially expressed between clusters were identified using ‘FindMarkers()’ function implemented in Seurat. Gene ontology (GO) enrichment analysis was performed using ToppGene Suite [44] and heatmaps were generated using custom R scripts. Single-cell pseudo-time trajectories of immune cell subpopulations were constructed with Monocle [45] as described before [46].

### Single-cell RNA sequencing of blood samples and data analysis

Single-cell suspensions of PBMCs were isolated from uninfected mice and mice at 6-days post-infection. Sequencing libraries were generated using Chromium Single-cell 3ʹ GEM, Library & Gel Bead Kit v3 (10x Genomics Cat # 1000075) on a 10× Genomics Chromium Controller following manufacturers’ protocol. Subsequently, libraries were sequenced on Illumina NextSeq 2000. Sequencing files were demultiplexed using “cellranger mkfastq”. Subsequently, “cellranger count” was used to perform alignment to mouse reference transcriptome (mm10), barcode processing and gene counting. Further analysis was performed using Seurat [43] as described above.

### Bulk RNAseq data analysis

We used publicly available data [20] to identify VEEV infection-induced changes in different regions of the brain. Raw counts were downloaded from Gene Expression Omnibus (GEO, accession ID: GSE213725). Data was then normalized using the Trimmed Mean of M-values (TMM) normalization method implemented in edgeR [47]. For differential expression analysis, the limma package [48] was employed in conjunction with the voom transformation [49]. Heatmaps for genes of interest were generated using the pheatmap package [50], and a boxplot was created using the ggplot2 package [51], both in R (version 4.3.1) [52].

### Flow cytometry

Cells were incubated for 30 min on ice in 100 μl Hank’s balanced salt solution (Thermo Fisher) + 2% FBS with Fc block (1:100 dilution, clone 2.4G2; BD Biosciences) along with the following antibodies, each diluted 1:500: CD45 APC-Cy7 (clone 30-F11; BD Biosciences), CD11b PE-CF594 (clone M1/70; BD Biosciences), CD3 PerCP-Cy5.5 (clone 17A2; BD Biosciences), CD4 BV650 (clone GK1.5; BD Biosciences), CD8 FITC (53-6.7; BioLegend), CD49b PE (clone DX5; BD Biosciences), and Perforin PE/Dazzle 594 (clone S16009A; BioLegend). Cells were then fixed using BD Cytofix/Cytoperm (BD Biosciences) according to manufacturer’s instructions. Flow cytometry was performed using a FACSAria Fusion and data were analyzed using FlowJo software. Microglia, other myeloid lineage, and lymphocytes were resolved using CD45 and CD11b expression, with microglia identified as CD45^int^ CD11b^int^, other myeloid as CD45^hi^ CD11b^hi^, and lymphocytes as CD45^hi^ CD11b^−^ as previously described [19].

### Cytokine analysis

Cytokines were quantified using LEGENDplex^TM^ multiplex bead-based assay (BioLegend) using the mouse anti-virus response panel according to manufacturer’s instructions. Flow cytometry of the beads was performed using a FACSAria Fusion and data were analyzed using BioLegend’s cloud-based analysis software available at https://legendplex.qognit.com.

### Immunohistochemical staining of mouse brains

Brains for immunohistochemical staining were collected from infected mice at 6 dpi. Upon euthanasia and perfusion as described above, mice were perfused with 20 mL 10% neutral buffered formalin (NBF). Isolated brains were subsequently immersed in 10% NBF at 4° C for 3 days with gentle agitation. Following fixation, brains were paraffin-embedded and cut into 5 μm thick sections. All sections were dewaxed with xylene and hydrated with alcohol. Citrate or Tris-EDTA were used for antigen retrieval, and hydrogen peroxide (ab64218, Abcam) was used to block endogenous peroxidase. After blocking non-specific sites with CAS-block (008120, Thermo Fisher Scientific), sections were incubated with Ly6c primary antibody (ab314120, Abcam) and secondary antibody (ab6720, Abcam). 3,3′-diaminobenzidine (DAB) kit (ab64238, Abcam) was used for visualization, and hematoxylin was used to stain the nuclei. All sections were rinsed with distilled water and sealed with permount (sp15-100, Fisher Scientific).

### Spatial transcriptomics

Spatial transcriptomic analysis of a VEEV-infected brain collected 6 days post-infection was conducted using 10x Visium technology (10x Genomics). 5 µm brain sections were mounted onto charged slides, deparaffinized, and stained with hematoxylin and eosin. The brain sections were imaged at 4x magnification using an ECHO Revolve microscope and the tiles were stitched together using Affinity Photo (Serif). The brain sections were then destained and decrosslinked to release RNA sequestered by formalin fixation. Next, the sections were incubated with mouse-specific probes targeting the whole transcriptome (PN-1000365, 10x Genomics), allowing for the hybridization and ligation of each probe pair. Gene expression probes were released from the tissue and captured by spatially barcoded oligonucleotides on the Visium slide surface using the Visium CytAssist instrument. Gene expression libraries were then prepared from each tissue section and sequenced using the Illumina NextSeq 2000.

After sequencing, the data and histology images were processed with the Space Ranger software (10x Genomics) and Seurat [43]. After demultiplexing the sequencing files with “spaceranger mkfastq”, the fastq files and the brightfield image of the tissue were provided to “spaceranger count” and read alignment, tissue detection, fiducial detection, and barcode/UMI counting were performed. Seurat was used then used for filtering (nFeature_Spatial > 500 & nCount_Spatial > 500 & percent.mt < 30), data normalization, dimensionality reduction, clustering, identification of spatially variable genes, and visualization. For each sample, after filtering, the data was normalized using “SCTransform()” function implemented in Seurat. Dimensionality reduction and clustering analysis was performed using “RunPCA()”, “FindNeighbors()”, “FindClusters ()” and RunUMAP() functions. We then applied the “FindAllMarkers()” function to identify genes enriched in each cluster. We also performed integrative analysis of uninfected and infected brain samples using the “SelectIntegrationFeatures()”, “PrepSCTIntegration()”, “FindIntegrationAnchors()”, and “IntegrateData ()” functions, followed by dimensionality reduction using PCA and UMAP. Clusters were visualized in UMAP space using DimPlot() and overlaid on the tissue image using “SpatialDimPlot()” function.

### Statistical analyses

Statistical significance was determined using Prism Version 10.2.3 (347) (GraphPad, La Jolla, CA). Specific tests and *p* values are indicated in the figure legends.

## Data Availability

The single cell RNA-sequencing and spatial data have been deposited at the NCBI Gene Expression Omnibus (GEO), accession numbers GSE274566 and GSE275201, respectively.

## Acknowledgements and Funding.

Funding for this research was provided by internal Lawrence Livermore National Laboratory Directed Research and Development funds (22-ERD-038 to D.R.W.). The funders had no role in study design, data collection and analysis, decision to publish, or preparation of the manuscript. This work was performed under the auspices of the U.S. Department of Energy by Lawrence Livermore National Security, LLC, Lawrence Livermore National Laboratory under Contract DE-AC52-07NA27344.

## Author Contributions

M.V.R., A.S., N.R.H, and D.R.W. conceived the project and designed experiments; M.V.R., N.F.L., A.M.P, N.R.H, and D.R.W. performed experiments; A.S. and B.M.G. analyzed sequencing data; D.R.W. supervised the project. M.V.R. and D.R.W. wrote the original draft, and all authors were involved in manuscript review and editing.

